# Transient thresholding: a mechanism enabling non-cooperative transcriptional circuitry to form a switch

**DOI:** 10.1101/134858

**Authors:** K. H. Aull, E. Tanner, M. Thomson, L. S. Weinberger

## Abstract

Threshold generation in fate-selection circuits is often achieved through deterministic bistability, which requires cooperativity (i.e., nonlinear activation) and associated hysteresis. However, the Tat positive-feedback loop that controls HIV’s fate decision between replication and proviral latency lacks self-cooperativity and deterministic bistability. Absent cooperativity, it is unclear how HIV can temporarily remain in an off state long enough for the kinetically slower epigenetic silencing mechanisms to act— expression fluctuations should rapidly trigger active positive feedback and replication, precluding establishment of latency. Here, using flow cytometry and single-cell imaging, we find that the Tat circuit exhibits a transient activation threshold. This threshold largely disappears after ∼40 hours—accounting for the lack of deterministic bistability—and promoter activation shortens the lifetime of this transient threshold. Continuous differential equation models do not recapitulate this phenomenon. However, chemical reaction (master equation) models where the transcriptional transactivator and promoter toggle between ‘inactive’ and ‘active’ states can recapitulate the phenomenon since they intrinsically create a single-molecule threshold transiently requiring excess molecules in the ‘inactive’ state to achieve at least one molecule (rather than a continuous fractional value) in the ‘active’ state. Given the widespread nature of promoter toggling and transcription factor modifications, transient thresholds may be a general feature of inducible promoters.

## INTRODUCTION

Thresholds allow biological systems to either respond to or disregard a signaling input, based on the input’s strength or level. Such thresholds are critical for cellular decision-making and are often a key design feature of gene-regulatory circuits, enabling the regulatory circuit to be robust to spurious signals or noise (1-3). Historically, the mechanism for threshold generation was thought to be either the presence of deterministic multistability (4-6) or zero-order ultrasensitivity (7, 8), both of which require specific regulatory architectures (high-order self-cooperativity with hysteresis and zero-order oppositional reactions, respectively). For example, if a putative activator molecule requires homo-dimerization (i.e., self-cooperativity) to become functional, this automatically generates a molecular threshold—determined by the dimerization disassociation constant—and can lead to deterministic bistability; below the dimerization threshold, there is no functional activator, whereas above the threshold, activation ensues.

Formally, deterministic multistability requires nonlinearity in the governing differential equations, which can be achieved by self-cooperative positive feedback:

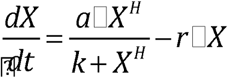

where X is the activator, *a* is the feedback strength, *k* is a Michaelis constant, *r* is the decay rate, and *H* is the Hill coefficient (Figure 1A, left). When the positive feedback is self-cooperative (i.e., *H* > 1), the circuit can exhibits deterministic multistability; in particular, if *H* = 2, the system can be bistable with two stable states (ON and OFF) separated by an unstable state, the ‘separatrix’. Bistable circuits exhibit a response threshold (specifically, at the unstable ‘separatrix’) and are characterized by hysteresis, a type of memory in which the circuit produces different dose-response curves, depending on whether signal increases or decreases (6). In contrast, positive-feedback circuits lacking self-cooperativity (*H* = 1) are monostable, having no separatrix (or threshold), no hysteresis, and only a single stable state; if this circuit can be turned ON, then the only stable state is the ON state (assuming the biochemical rate constants are not changing), with the OFF state being necessarily unstable (Fig. 1A, right).

**Figure 1:**
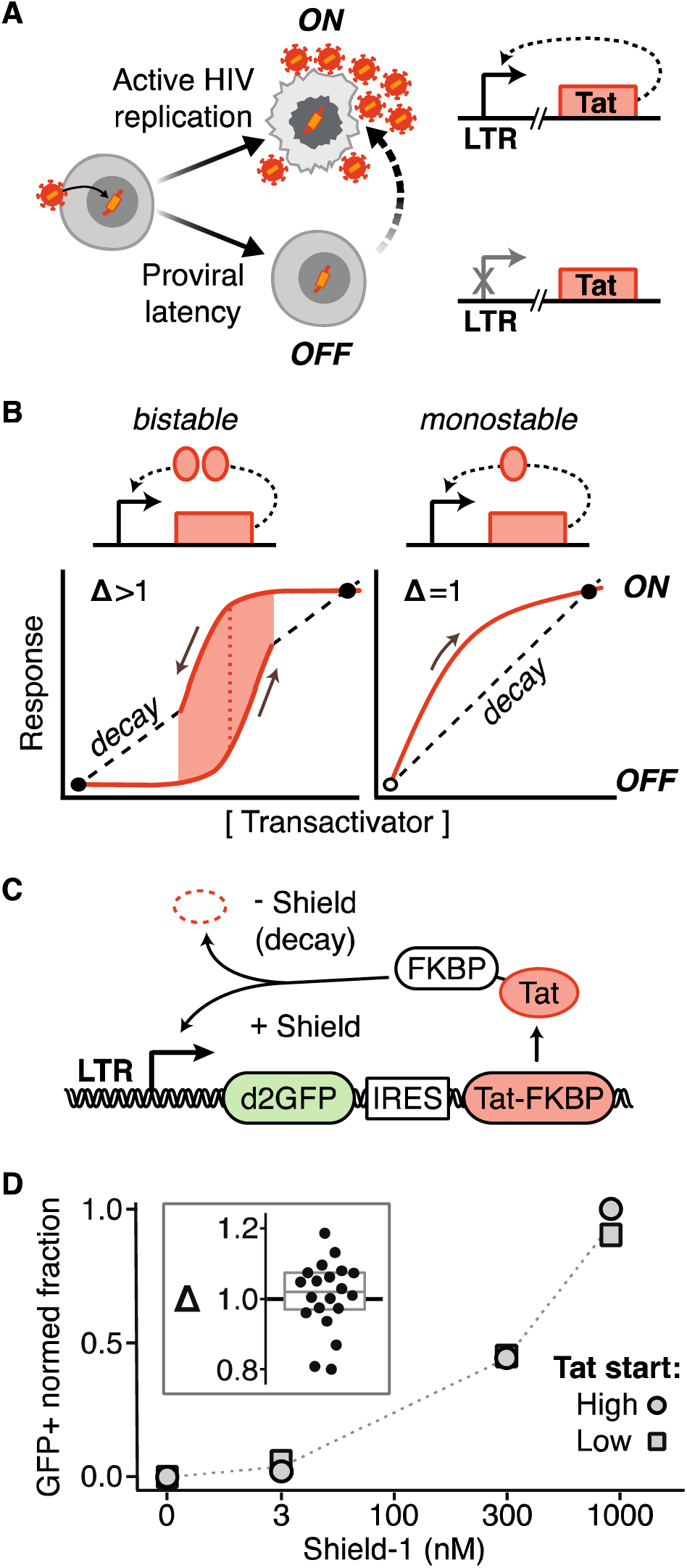
The HIV LTR-Tat positive-feedback circuit lacks hysteresis and bistability. (A) Schematic of the HIV fate decision between active replication (ON) and latent (OFF) states. This fate decision is controlled by the HIV Tat-LTR positive-feedback circuit. Transactivation of LTR, the sole promoter of HIV, by its gene product Tat drives further Tat production and HIV replication. (B) Bistability versus monostability in positive-feedback transcriptional circuits. Formally, deterministic multistability requires nonlinearity in the governing differential equations (6); for example, if the activator requires cooperative self-association to bind its promoter, its expression is described by a nonlinear Hill equation (Hill coefficient *H* > 1) (20). Such circuits exhibit bistability, having two attractor states (ON and OFF) separated by a response threshold—at low activator levels, the decay rate (dashed line) dominates over synthesis (solid line), and at high levels, the opposite is true—and hysteresis, a type of memory in which the response is history dependent, following different paths from ON-to-OFF versus OFF-to-ON (the difference between paths is Δ > 1). In contrast, circuits lacking self-cooperativity (*H* = 1) are monostable, having neither a threshold nor hysteresis (Δ = 1)—if a monostable circuit can be turned ON, its only stable state is the ON state (assuming the biochemical rate constants are not changing), with the OFF state being necessarily unstable (6). (C) Schematic of the minimal HIV “Ld2GITF” positive-feedback circuit used to test for hysteresis (<u>L</u>TR driving a <u>2</u>-hour half-life <u>GFP</u> reporter and an IRES expressing <u>T</u>at fused to <u>F</u>KBP, a degradation tag inactivated by the small molecule Shield-1). (D) Hysteresis test by flow cytometry analysis of Ld2GITF. Isoclonal Jurkat Ld2GITF cells were either pretreated with 1 μM Shield-1 for 4 days to activate cells to start in an ON start (oval data points) or not pretreated to start in an OFF state (square data points). All cells were washed and then incubated in the specified Shield-1 alongside for an additional 4 days, and the percentage of GFP+ cells was measured. Inset: Δ (the ratio of pretreated to not-pretreated GFP+ cells) calculated for five isoclonal populations of Ld2GITF (<Δ> ≈ 1).

Gene-regulatory circuits typically achieve *H* > 1 and bistability via cooperative binding of a transcription factor to its promoter (9, 10). Notable examples of bistable gene-regulatory circuits include the toggle switch (11), phage lambda lysis-lyogeny (12, 13), the lac operon (14), and competence in *Bacillus subtilis* (15, 16), all of which have thresholds established by high-cooperativity feedback loops. Other mechanisms for generating a threshold include zero-order ultrasensitivity (7, 8) and buffered threshold-linear responses (17, 18); however, when applied to transcription-factor induction of a promoter, these models (19) either fail to generate a threshold response (see Appendix 1) or rely on an excess of substrate (i.e., the promoter itself) (20), respectively.

In stark contrast to these canonical examples, the circuit that controls HIV’s fate decision between active replication and proviral latency (Fig. 1B) appears to lack the classic mechanisms associated with deterministic bistability or ultrasensitivity (21). Latent HIV is the chief barrier to a cure (22) and the decision between active replication and latency in HIV is governed primarily by the virus’s positive-feedback circuit in which HIV Tat protein transactivates expression of the HIV long terminal repeat (LTR) promoter, the only promoter in the virus (Fig. 1B). During latency, the LTR is largely quiescent but establishment of latency is not correlated with viral integration site (23-25) or progressive cellular silencing (26). Specifically, epigenetic silencing occurs on the order of weeks (27), whereas ∼50% of infections result in *immediate* establishment of latency *in vitro* (28, 29), and latency is established within 72 hours *in vivo* (30). Overall, latency *establishment* occurs too quickly to be accounted for by epigenetic silencing, which acts on timescales of weeks in T cells (27). Instead, the data appear more parsimonious with the Tat-LTR positive-feedback circuit being necessary and sufficient for establishment of latency (26), while long-term stability of latency is likely mediated by epigenetic silencing (31).

Tat acts as a monomeric transactivator, binding to a single site on a nascent RNA hairpin formed by stalled RNA polymerase II at the LTR promoter. Because Tat binds non-cooperatively, classical deterministic models predict that the circuit should have no activation threshold and thus the latent state would be unstable (32). Thus, it is unclear how, without bistability, HIV generates a molecular threshold in Tat so that it can even temporarily remain in an off state and provide an opportunity for the kinetically slower epigenetic silencing mechanism to act. Given the noisy expression of the HIV LTR promoter (33, 34), Tat positive feedback should trigger active replication within these first few days. This would preclude establishment of proviral latency, as active replication destroys the cell within hours (35) and silencing of an actively replicating cell cannot overcome active HIV gene expression (26). In general, it remains unclear how the Tat positive-feedback circuit that lacks deterministic bistability (and ultrasensitivity) can generate a threshold to establish a stable off state.

Here, we examine the HIV Tat-LTR circuit to determine how a threshold can be generated without self-cooperativity. Using a combination of single-cell experimental analyses, both flow cytometry and time-lapse fluorescence microscopy, we find that the LTR circuit exhibits a transient threshold for activation by Tat. The threshold gradually disappears, and at ∼40 hours, there appears to be no effective threshold such that the LTR-Tat circuit exhibits no hysteresis or deterministic bistability and cellular activation (e.g., NF-κB signaling), which modulates the kinetics of promoter toggling, shortens the transient lifetime of the threshold. Stochastic model where the transcriptional transactivator and promoter toggle between ‘inactive’ and ‘active’ states appear sufficient to recapitulate the transient threshold phenomenon.

## MATERIALS AND METHODS

### Cell lines and reagents

The minimal Ld2GITF feedback circuit and the doxycycline-inducible Tat-Dendra cell line have been described (26). Here, the lentiviral LTR mCherry reporter from (26) was modified to contain an N-terminal PEST tag, giving LTR mCherry-deg, with mCherry protein half-life 10.7 hours (data not shown). Plasmid maps and cloning details available on request. LTR mCherry-deg was packaged in 293T cells and used to infect Jurkat Tat-Dendra cells at low MOI (mCherry positive cells < 5%). These cells were induced at high Dox (500 ng/mL) for 2 days, and FACS sorted to isolate dual-positive single cells with a FACSAria II (BD Biosciences USA, San Jose, CA) that were grown into isoclonal populations. Isoclones were screened to confirm robust dual-positive response to Dox with negligible expression at baseline. Unless otherwise stated, all chemical reagents were sourced from Sigma-Aldrich USA (Saint Louis, MO). When specified, the HIV reactivating agents TNF (10 ng/mL tumor necrosis factor alpha) or TSA (400 nM trichostatin A) were supplied at the time of Dox addition.

### Flow cytometry data collection and analysis

To generate dose-response plots, each isoclone and condition was tested at eight doxycycline (Dox) levels: seven twofold dilutions, from 250 to 3.9 ng/mL, plus a zero-Dox control. Data were collected on a MACSQuant™ high-throughput flow cytometer (Miltenyi Biotec, Bergisch Gladbach, Germany), gated for live single cells in FlowJo™ (Tree Star, Ashland, OR). The mCherry positive cutoff was chosen to exclude non-induced cells. All eight Dox dilutions were pooled and cells were grouped by Tat-Dendra signal to estimate the conditional probability of LTR response for the specific Tat level. A schematic of this workflow, with sample data, is presented in Fig. 2B-D (all Tat-Dendra values were background-subtracted, using the mean of zero-Dox control as background; clusters with non-positive Tat-Dendra values were not considered in the analysis). The dose-response and dose-mean expression curves obtained by this method were fit to a standard Hill function: 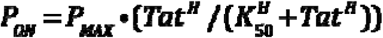. Nonlinear least-squares fitting was performed in R, using the nlsLM function from the minpack.lm package (CRAN).

**Figure 2:**
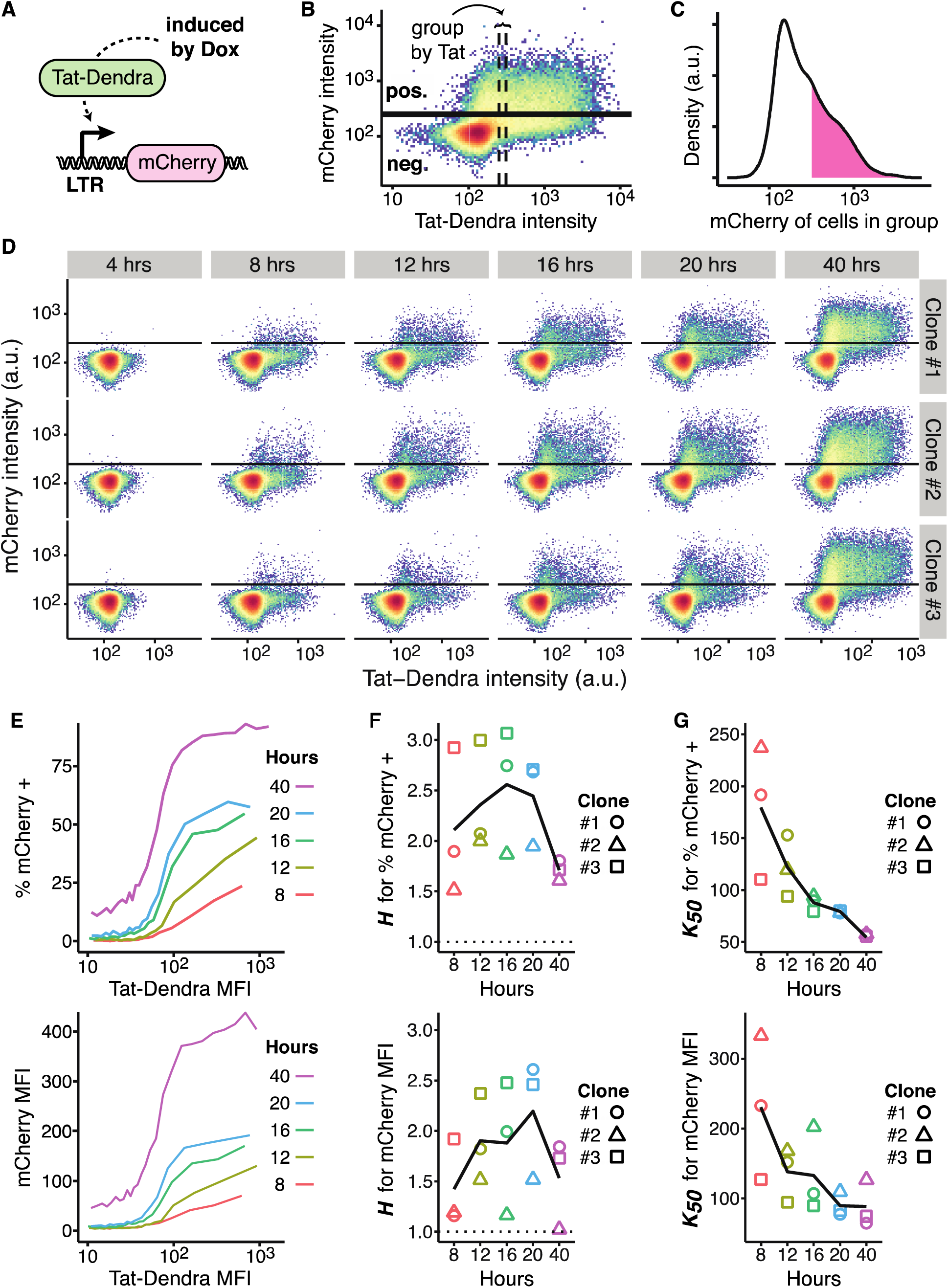
The LTR promoter exhibits a transient threshold in its response to Tat. (A) Schematic of constructs used to directly quantify LTR-Tat dose-response function. A doxycycline-inducible promoter (Tet-ON system) drives expression of a Tat-Dendra2 fusion protein, which activates the LTR promoter to drive expression of destabilized mCherry reporter. (B) Scheme for estimating conditional probability of LTR activity for a given Tat level, with a representative two-color flow cytometry dot plot of an isoclonal Jurkat cell line stably expressing constructs in panel A after 20 hours of Dox induction (plot shown is Clone #2). To estimate the conditional probability of LTR expression for a given Tat level, data was combined from eight Dox dilutions (0–250 nM). The dense spot in the lower left corner corresponds to non-induced cells (i.e., auto-fluorescence background), which the mCherry positive cutoff gate excludes (as indicated by the black horizontal line). Cells with similar Tat-Dendra values were grouped, as indicated by the vertical dashed lines, and the percentage of cells above the mCherry positive cutoff and mean mCherry fluorescence was recorded for each group. For visual clarity, this panel depicts a group of 2500 cells, while the analysis uses a tighter group of 1000 cells. (C) Histogram of mCherry intensity for cells in the marked group. Density above the mCherry positive cutoff is shaded. Despite the narrow band of Tat-Dendra intensities, the LTR response is variable. (D) Full flow cytometry time-course for three isoclones of Jurkat encoding both Tet-Tat-Dendra and LTR-mCherry-deg induced with eight Dox dilutions, and measured by flow cytometry over time. Horizontal lines indicate the mCherry positive cutoff. At early times, a pronounced “shoulder” is visible in Tat expression where a substantial percentage of cells express Tat-Dendra but these cells do not express mCherry from the LTR. (E) Calculated dose-response curves for percentage of mCherry+ cells (top) and mCherry mean fluorescence intensity (MFI, bottom) from data in panel D. Clone #1 is shown; the other isoclones, and Hill fits, are presented in Fig. S1 (Supporting Materials). (F) Calculated Hill coefficients (*H*) from dose-response curves over time. The expected non-cooperative response (*H* = 1) is indicated by a dashed line, all data points are above the expected H = 1 line. Maximum *H*-values occur at intermediate time points for both %mCherry cells and MFI. (G) Calculated half-maximal response (K_50_) from fits of the dose-response curves over time. K_50_ declines over time, indicating that the threshold becomes progressively weaker.

### Immobilization of cells for time-lapse imaging

5–10x10^6^ actively dividing (healthy) Jurkat cells were washed twice in regular phosphate buffered saline (PBS), then again in mildly alkalized PBS (pH 8.0). Immediately before use, a single aliquot of biotinylation reagent (1 mg EZ-Link Sulfo-NHS-LC-Biotin, ThermoFisher, Waltham, MA) was re-suspended in 800 μL PBS (pH 8.0). Of this, 500 μL was used to re-suspend the cells after the final wash, while the rest was added to a collagen-coated coverslip plate (#1.5, 35 mm, MatTek, Ashland, MA). Both cells and coverslip were kept at room temperature. After 30 min, the coverslip was thoroughly rinsed with PBS + 50 mM glycine, then coated with 80 μL streptavidin (1 mg/mL, New England Biolabs, Ipswich, MA). The cells were washed twice in glycine solution, then again in standard culture medium. During the final wash step (∼15 min later), the coverslip was rinsed with PBS to remove unbound streptavidin. The biotinylated cells were resuspended in ∼300 μL culture medium, transferred to the coverslip, then placed in the incubator for 30 min to settle by gravity. Unbound cells were then carefully rinsed away, and the plate was refilled with 2.5 mL of culture medium containing 250 ng/mL Dox. The finished plate was placed on the microscope for thermal equilibration (∼1 hr) and subsequent imaging.

### Microscope setup and imaging conditions

All imaging was performed on a Zeiss AxioVert inverted fluorescence microscope (Carl Zeiss, Jena, Germany), equipped with a Yokogawa spinning disc, CoolSNAP HQ2 14-bit camera (Photometrics, Tucson, AZ), and laser lines for 488 nm and 561 nm excitation. To facilitate time-lapse imaging, the microscope has a programmable stage with Definite Focus, and also a stage enclosure that maintains samples at 37°C and 5% CO_2_ with humidity. Images were captured every 10 min, sampling a 5x5 X-Y grid, one Z-position each. Exposures were 800 ms at 20% power with the 561-nm laser, then 400 ms at 10% power with the 488-nm laser, then 600 ms for brightfield. The objective used was a 40X oil, 1.3NA, with 2x2 camera binning applied. For all “induced Tat” movies, imaging was started no more than 2.5 hours after Dox addition, and was continued until 20 hours. For protein half-life measurements, imaging was started 10 min after addition of 10 μg/mL cycloheximide and continued for 50 10-min intervals. Bleaching half-life was measured with the same image settings, but taken at one location in 5-sec intervals to minimize changes in total protein level. For HSV-GFP imaging, to maximize the visibility of these very small particles, the 488-nm exposure time was increased to 40 sec and binning was turned off. For each location in a 7x7 X-Y grid, nine Z-positions were sampled at 0.2 μm intervals; the most in-focus image was chosen for analysis.

### Image segmentation analysis to generate single-cell trajectories

The center of each cell was manually marked, using the final brightfield image and a custom script (MATLAB, The MathWorks, Natick, MA). For each cell location, a 23-pixel diameter circle was marked around it, and the mean fluorescence intensity within that circle was recorded at each time point to generate single-cell trajectories. Each trajectory was then subjected to automated quality control (QC): cells in which any two consecutive readings differed by more than 15% in either channel were excluded; upon review of the source images, these events were typically due to cell division, or another cell drifting into view. Cells that began the experiment “on” were also excluded (LTR >2% over background at 2.5 hours post-Dox addition; this was rare, 2–5 cells per condition). Illustrations of the raw image data and QC process are available in Fig. S2 (Supporting Materials). For these movies, between 2001 and 2193 cell trajectories passed QC. The trajectories were normalized to set their lowest values to zero, then fit to a smoothing spline in base R (df=10, n=105) to further reduce noise. Tat-Dendra trajectories were also corrected for photobleaching. This was not necessary for mCherry, which did not bleach under the imaging conditions used (data not shown). The photobleaching correction process is described in Fig. S3 (Supporting Materials).

### Quantitation of Tat-Dendra molecular number by GFP molecular rulers

For quantitation using the HSV-GFP molecular ruler (36, 37), the images of viral particles were processed using a custom script (MATLAB, The MathWorks, Natick, MA). Each image was background-subtracted, using the median of all 49 images as background, then thresholded to include the bright particles and the first Airy disk surrounding them. The MATLAB function bwconncomp was used to identify potential features within the images. To set the correct size, TetraSpeck™ beads (ThermoFisher, Waltham, MA) were analyzed by the same method; the 0.2- μm beads were 15–18 pixels (data not shown). Since the HSV-1 capsid is 125 nm (38), features between 10–14 pixels were selected. For each feature, the total intensity above background was recorded. The mean value was 1424 units. (95% CI 1412–1435; n=5004.). Given that the HSV-GFP images had 100X the exposure time, and 4X as many pixels, relative to the Tat-Dendra images, each intensity unit of HSV-GFP represents 25X less signal. EGFP is also brighter than Dendra2 by 1.47X (39) such that there are [1424 / 25] intensity equivalents per [900 x 1.47] molecular equivalents, which reduces to 1 intensity unit per 23.2 Tat-Dendra (Fig. S4, Supporting Materials). From the single-cell imaging data, the threshold level of Tat proteins required to minimally activate the LTR (i.e., > 2% mCherry positive cells) gives an intensity signal of 5.0 units per pixel, or 1900 units per cell (each cell is 377 pixels). The conversion factor calculated from molecular ruler thus estimates the minimal activation threshold at 4.4x10^4^ Tat molecules per cell (Fig. 3C).

**Figure 3:**
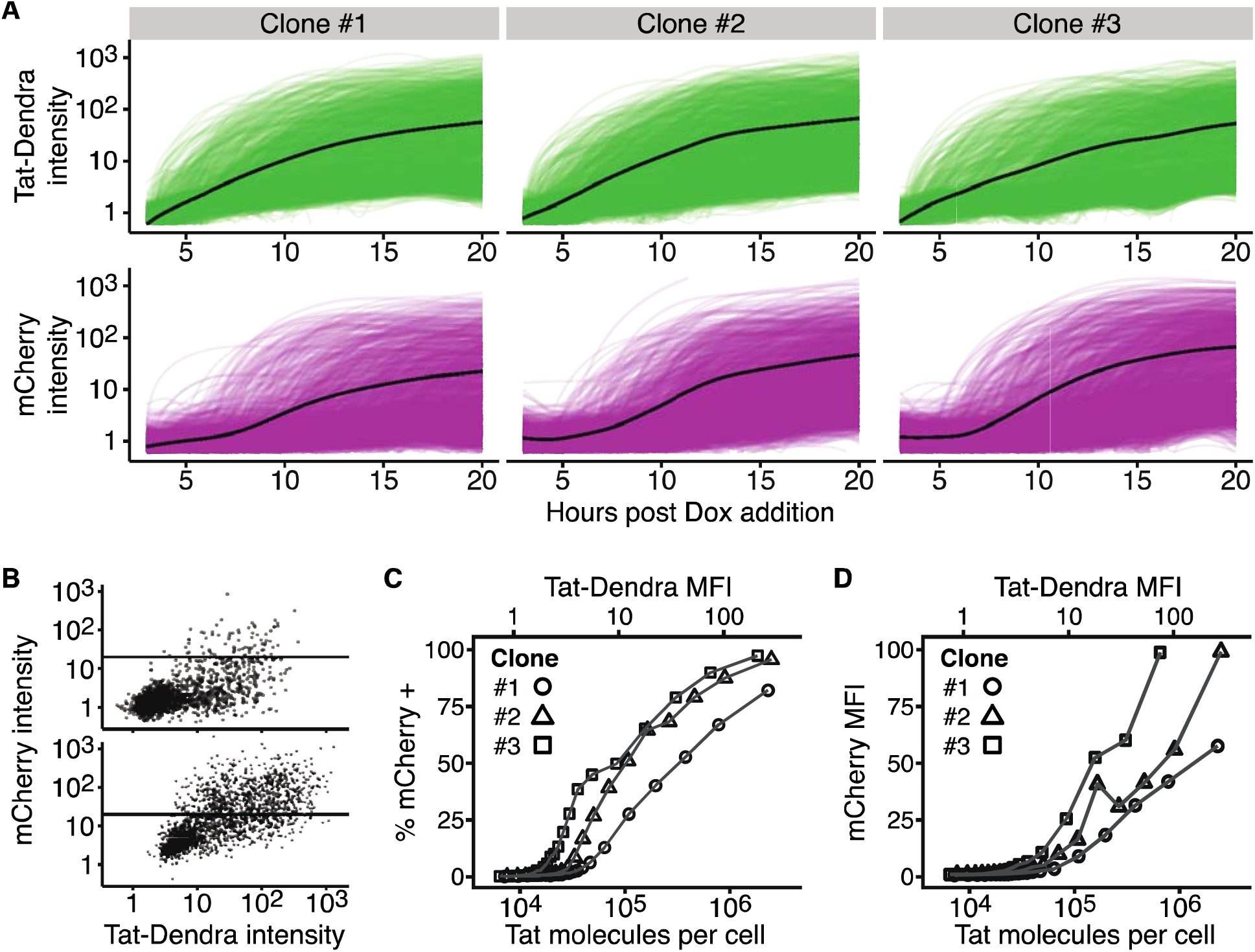
Time-lapse microscopy verifies that the LTR exhibits an activation threshold at early times. (A) Time-lapse fluorescence microscopy imaging of single cells from three Jurkat cell isoclonal populations each encoding both Tat-Dendra and LTR-mCherry-deg. Cells were activated and then imaged for 20 hours. Tat-Dendra trajectories are green; mCherry trajectories are magenta; mean intensity trace shown in black. ∼2000 cell trajectories shown for each clone. (B) Flow-style dot plot of Tat-Dendra versus mCherry intensities from time-lapse images at t = 10 h (upper) and t = 20 h (lower) of Clone #2. Each dot represents an individual cell. As in Fig. 2, the horizontal line marks the mCherry-positive cutoff gate. (C-D) Dose-response curves for %mCherry+ cells (left) and mCherry mean fluorescence intensity (MFI, right) versus Tat MFI and calculated number of Tat molecules per cell. Single-cell intensities extracted from all images were pooled and processed in the same manner as the flow data (each point summarizes 10^4^ observations). Tat-Dendra signal intensity was converted to molecular number using a GFP “molecular ruler”.

### Computational modeling

Deterministic and stochastic computational modeling (Appendix I and Fig. 5, respectively) was carried out in Mathematica™. Deterministic ordinary differential equation (ODE) models of Tat transactivation of the LTR were based on generalized mathematical models of ultrasensitive responses (19) and previous experimentally validated LTR-Tat circuit models (32) that incorporate reversible acetylation-deacetylation of Tat protein (i.e., so-called ‘futile cycles’). For stochastic models of chemical master equations, the two-state model of the LTR promoter (40-43) was simulated by Gillespie’s method (44) using the Mathematica™ xSSA package (http://www.xlr8r.info/SSA/). The outputs from simulations are presented in arbitrary numbers. Initial conditions for all species were set to 0 (except LTR_OFF_ = 1) and simulations were run to time=200 (arbitrary time units, 200 simulations were run per model and parameter set, and mean Tat and mCherry values for all runs were calculated at specified time points.

## RESULTS

### The HIV LTR-Tat circuit lacks hysteresis and bistability

Previous studies demonstrated that the HIV Tat-LTR positive-feedback loop exhibits a purely linear expression rate at early times (i.e., scales linearly with Tat and lacks cooperativity) (32), as expected for non-cooperative positive feedback (Fig. 1A). To confirm that the LTR-Tat circuit does not establish bistability through other mechanisms (e.g., nonlinear degradation), we tested for hysteresis in a minimal Tat-LTR feedback circuit, where LTR drives expression of an unstable (2 hr half-life) GFP reporter (d2GFP) and an IRES enables co-expression of Tat fused to the tunable proteolysis tag FKBP (Fig. 1C). In this circuit (hereafter Ld2GITF), Tat proteolysis can be protected by the small molecule Shield-1 (45), thereby allowing feedback strength to be tuned (26) and alternate paths of the circuit—ON-to-OFF versus OFF-to-ON—to be examined. Specifically, cells in the GFP ON state (i.e., pre-incubated in Shield-1) can be exposed to successively decreasing Shield-1 levels to examine turning OFF of the circuit, while cells in the GFP OFF state (i.e., no Shield-1 pre-incubation) can be exposed to successively increasing Shield-1 levels to examine turning ON of the circuit. The difference (Δ) in percentage of GFP ON cells for a specific Shield-1 concentration can be quantified, with Δ > 1 indicating hysteresis. If hysteresis is present, cells beginning in the ON state (i.e., pretreated with high Shield-1) will be more likely remain ON at a specific intermediate dose of Shield-1, as compared to cells that began in the OFF state (i.e., non-pretreated cells); whereas, if hysteresis is not present (Δ = 1), there will no difference in ON-OFF percentages for cells beginning in either the ON or OFF state. We tested five isoclonal Ld2GITF populations carrying single integrations of the Ld2GITF circuit, and measured Δ to be ≈1 (Mean 1.008; 95% CI 0.813-1.203; Fig. 1D), indicating that hysteresis is unlikely. These hysteresis measurements build upon previous data indicating that the necessary conditions for deterministic bistability are absent in the HIV Tat-LTR circuit (32).

### Single-cell flow cytometry analysis of the HIV LTR-Tat dose-response function shows a threshold-like response that is transient in time

Absent bistability, it was unclear how the Tat-LTR circuit might encode a threshold to temporarily remain OFF to provide an opportunity for the kinetically slower epigenetic-silencing mechanisms to act. Importantly, chromatin-silencing mechanisms appear unable to silence the actively transcribing promoter (26).

First, to check if the Tat-LTR circuit encodes an activation threshold, we directly quantified LTR activity as a function of Tat levels using an ‘open-loop’ Tat-LTR dose-response system. In this system, one construct encodes Tat fused to the fluorescent reporter Dendra2 expressed from a doxycycline-inducible tet promoter, while a second construct encodes an mCherry reporter expressed from the LTR promoter (Fig. 2A). This open-loop system allows Tat levels to be tuned by doxycycline (Dox) and enables both Tat (dose) levels and LTR (response) levels to be quantified in the same cell (26) so that the dose-response ‘transfer’ function for Tat and LTR can be fit and an effective Hill coefficient calculated.

To estimate the conditional probability of LTR mean expression level and percentage ON for a given Tat level from flow cytometry data, a binning method similar to previous methods (46) was used (Fig. 2B-C). Examination of the flow cytometry time-course data showed that the LTR appears essentially non-responsive to Tat at low Tat levels, but LTR activity then increases sharply over a narrow range of Tat (Fig. 2D). At early times after Dox activation, a pronounced “shoulder” is visible in Tat expression where a substantial percentage of cells express Tat-Dendra, but these cells do not express mCherry from the LTR. This delay between Tat-Dendra and mCherry expression is on the order of 8–12 hours, which is too long to simply be a temporal delay in expression of mCherry due to activation by Tat-Dendra.

For all LTR isoclones (i.e., integration sites) examined, the dose-response expression curves for mCherry mean expression and percentage of mCherry ON cells exhibit a conspicuous activation threshold (Fig. 2E). The LTR appears essentially non-responsive to Tat at low Tat levels, but LTR activity then increases sharply over a narrow range of Tat. This thresholding behavior appears to be maximized at intermediate time points of 16–20 hours (Fig. 2F-G). At early times, the response is incomplete, but by 40 hours, the dose-response curves flatten with the K_50_ shifting to lower Tat expression.

### Time-lapse microscopy analysis verifies the threshold-like LTR response to Tat at early times after activation

To verify that this result was not simply a peculiarity of the flow cytometry approach, we next examined activation of this ‘open-loop’ activation circuit using quantitative time-lapse imaging (Fig. 3A). Jurkat isoclones, as above, were imaged for 20 hours after Dox activation, and for all isoclones, there was a conspicuous delay of approximately seven hours in mCherry expression relative to Tat-Dendra expression (Fig. 3A–B). The single-cell trajectories were then used to construct Tat-LTR dose-response trajectories via the same conditional binning method as used for flow cytometry (Supporting Material). For all LTR isoclones examined, the dose-response expression curves for both mCherry mean expression and percentage of mCherry ON cells exhibits a conspicuous activation threshold (Fig. 3C–D). As observed in flow cytometry, the microscopy imaging shows that the LTR is essentially non-responsive to Tat at low Tat levels, but LTR activity then increases sharply over a narrow range of Tat.

We used a ‘molecular ruler’ approach (36, 37) to convert Tat-Dendra fluorescence levels to molecular number (Methods and Supporting Material). For all clones tested, the threshold level of Tat proteins required to minimally activate the LTR (i.e., > 2% mCherry positive cells) is in the tens of thousands of molecules, with the average being 4.4x10^4^ Tat/cell (Fig. 3C–D). Comparable values for Tat molecules per cell were previously obtained in a minimal Tat-LTR feedback circuit, with quantitation performed by GFP standard beads (47). Upon accounting for cell size differences, this molecular threshold value was also not dissimilar to those calculated for phage lambda, where 55 Cro molecules are required for lytic infection and 145 CI molecules are required for lysogeny (13); human lymphocytes are ∼10^3^ times the volume of *E. coli* (48).

### Transcriptional activation by TNF effectively accelerates the transient lifetime of the LTR activation threshold

Based on observations that HIV latency can be partially reversed by transcriptional activators, we next asked if transcriptional activators could alter the observed LTR-activation threshold. To transcriptionally active the LTR, we used the well-characterized cytokine tumor necrosis factor alpha (TNF), which acts through nuclear factor kappa B (NF-κB) signaling to recruit transcriptional activators to the LTR (26, 49, 50), thereby increasing LTR transcriptional burst frequency (33, 51, 52).

When the dose-response function is measured post Dox induction in the presence of TNF, the response functions show a both marked shortening of the lifetime of the threshold and a reduced threshold (Fig. 4A and Fig. S5–6, Supporting Materials). In fact, when comparing the dose-responses in the presence and absence of TNF, the presence of TNF caused the 20-hour dose-response curve to look similar to the 40-hour non-TNF dose-response curves (compare Fig. 4A to Fig. 2E). Consistent with this observation, the calculated Hill coefficients, *H*, decreases in the presence of TNF (Fig. 4B) and, with the exception of clone #2 MFI, the K_50_ values decline in the presence of TNF (Fig. 4C), indicating that the threshold becomes progressively weaker.

**Figure 4:**
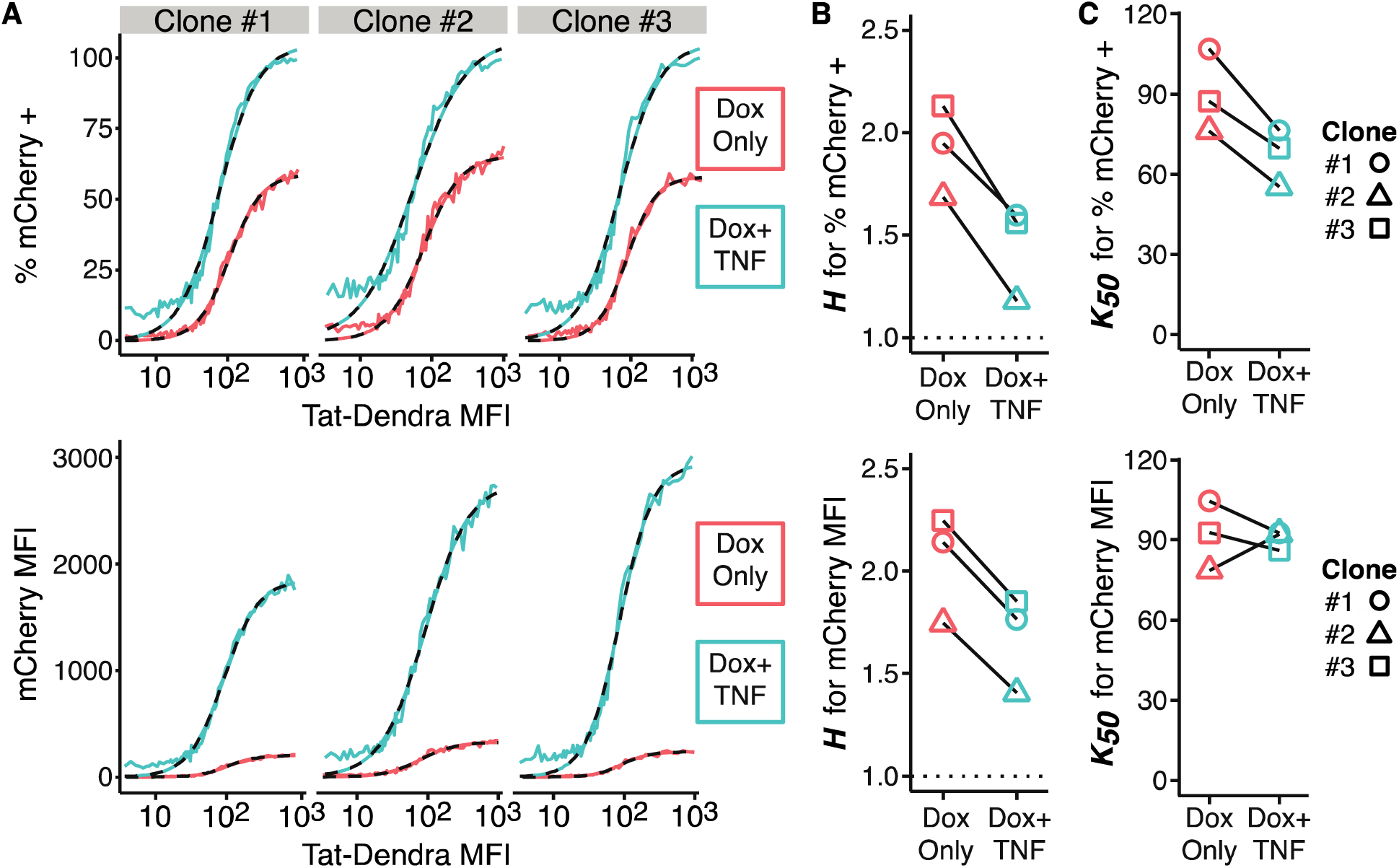
Transcriptional activation by TNF effectively accelerates the transient lifetime of the LTR activation threshold. (A) Dose-response curves for %mCherry-positive cells (top) and mCherry MFI (bottom) from flow cytometry measurements of three isoclones of Jurkat Tet-Tat-Dendra + LTR-mCherry, at 20 hours post Dox induction in the presence or absence of TNF. Each data point depicts a group of 500 cells. These data were fit to a Hill function (dashed lines); numeric results are given in Fig. S6 (Supporting Materials). (B) Hill coefficients, *H*, determined from dose-response curve fitting, demonstrate empirical positive cooperativity (*H* > 1) with lowering of *H*-values for cells treated with TNF. The expected non-cooperative response is indicated by the dotted line at *H = 1*. (C) Half-maximal response (K_50_), determined from dose-response curve fitting.

### A minimal stochastic model is sufficient to recapitulate the transient-threshold effect

We next explored whether a mechanistic model could be developed to explain the transient threshold effect. Given the lack of bistability (32) and hysteresis in the circuit (Fig. 1), we neglected models that postulated built-in cooperative responses or deterministic thresholds (i.e., models with a deterministic *H* >1).

Based on previous literature on ‘ultrasensitive’ threshold responses (19), we first examined a set of deterministic ordinary differential equation (ODE) models (Appendix I) that are non-cooperative (i.e., *H* = 1) but have architectures found in ultrasensitive responses, namely enzymatic inter-conversions in the zero-order regime. In these models, Tat is reversibly covalently modified—acetylated at lysine residues by p300 and de-acetylated by SirT1, with acetylation required for efficient transactivation of the LTR but deacetylation being more rapid than acetylation (32, 53). The rationale for testing these models was that the Tat-Dendra reporter (Fig. 2) does not distinguish between acetylated and deacteylated Tat and most Tat in the cell is deacetylated (32, 53), so Dendra intensity primarily quantifies Tat that is not transactivating the LTR. Moreover, given the faster deacetylation rate, a large amount of deacetylated Tat protein is required for significant acetylated Tat to be present. Nevertheless, in the deterministic regime, models of this form do not generate threshold responses either at steady state or in the pre-steady-state transient regime (Appendix I). This is because—without postulating an *ad-hoc* threshold for Tat acetylation—the continuous nature of deterministic ODEs results in a small fractional value of Tat protein continuously acetylated and thus transactivation competent.

Given the continuous nature of ODE models, we next examined minimal stochastic chemical reaction (master equation) models as these models account for integer molecule numbers. These models intrinsically form a threshold since a single molecule of active transactivator (rather than a continuous fractional value) is required for a reaction to occur. As above, we hypothesized that the rates of conversion from the ‘inactive’ to the ‘active’ state, could allow many transactivator molecules to be transiently present in the ‘inactive’ state before a single molecule of active transactivator (acetylated Tat) is produced, thereby establishing a transient threshold. To test this hypothesis, four stochastic models of increasing complexity were built (*models i–iv*, below).

The models are presented using a generic nomenclature where the active ‘transcription factor’ (TF) represents Tat; the TF can be in an inactive form (TF_i_) requiring a single modification to become active TF or (TF_ii_) requiring two modifications to become active TF. The promoter, which represents the LTR, can toggle between an ‘on’ state (Pr_on_) or ‘off’ state (Pr_off_). For computational expediency and simplicity, the models are course grained to neglect the mRNA intermediate.

where all models (*i–iv*) also include the following common reactions:

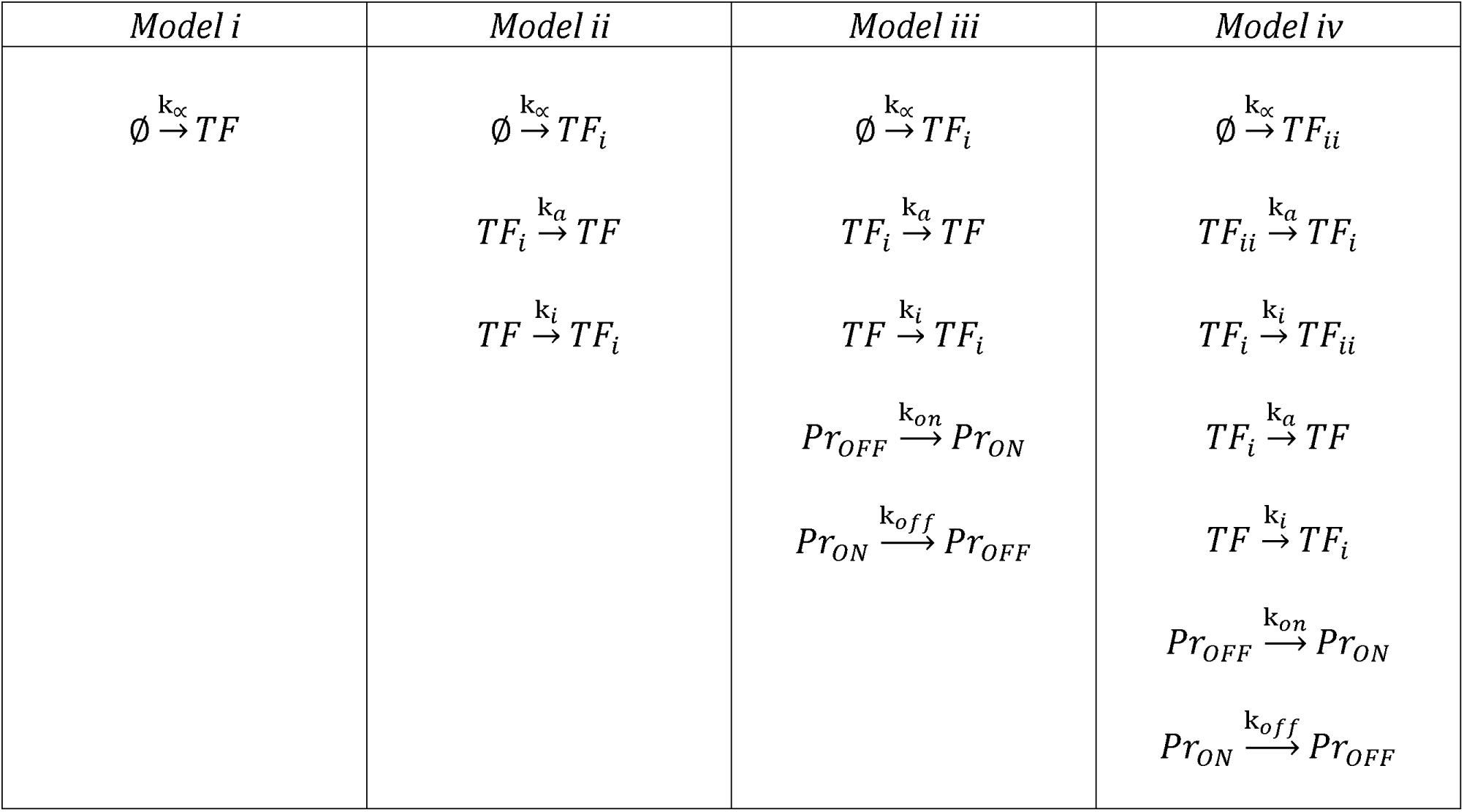

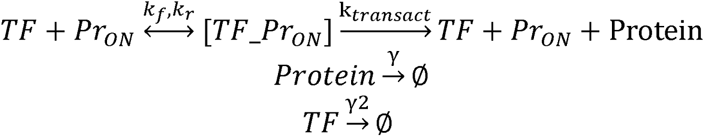

Models *i–iv* were then numerically simulated for the LTR-Tat system [*TF* = Tat-Dendra, *Pr* = LTR, *Protein* = mCherry, and k_α_ is Dox induction; (Fig. 5)]. In *model i,* active *TF* is produced at linear rate k_α_, transactivates *Pr*_*on*_ by forming the [TF_Pr_on_] complex and, as expected, generates linear dose-responses for *Protein* (mCherry) as a function of *TF* (Tat-Dendra) (Fig. 5). When the model is extended (*model ii*) so that *TF* is produced as inactive and reversibly modified to active (*TF*_*i*_↔*TF*), a slight threshold in dose response appears at early times (Fig. 5). The lifetime of this transient threshold is extended by inclusion of promoter toggling (*model iii*) and further extended (*model iv*) by additional transactivator toggling reactions (*TF*_*ii*_↔*TF*_*i*_↔*TF*) (Fig. 5).

**Figure 5:**
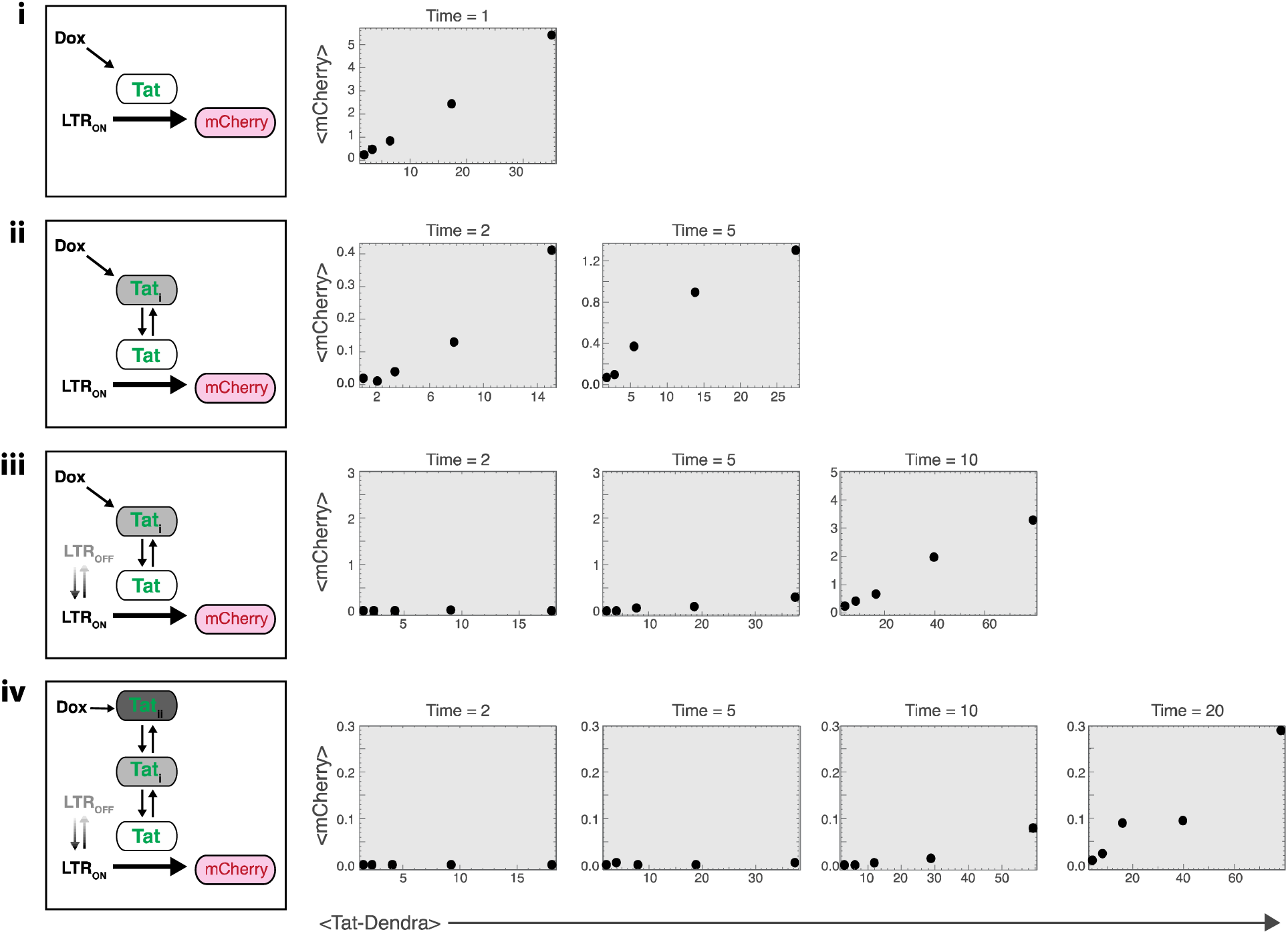
Stochastic models where the transactivator and promoter toggle between ‘active’ and ‘inactive’ qualitatively recapitulate the transient dose-response threshold. (Left) Schematics of models *i-iv* for the LTR-Tat system (i.e., *TF* = *Tat*, *Pr* = *LTR*, *Protein* = *mCherry*, and *k*_*a*_ is *Dox* induction). In all models, Tat is generated from the Dox-inducible promoter (at rate k_α_), and mCherry is driven from the LTR. (Right) Corresponding dose-response functions from stochastic simulations. Mean mCherry and mean Tat-Dendra calculated from 200 simulation runs at each specified time point (arbitrary time units). Both active transactivator (*Tat*) and inactive transactivator (*Tat*_*i*_ and *Tat*_*ii*_) are Dendra labeled; reverse reactions (e.g., LTR_OFF_→LTR_ON_ and Tat→Tat_i_) are 10-fold faster than forward reaction rates. (i) A model where neither the promoter toggles (only LTR_ON_) nor the transactivator toggles (only Tat) exhibits a linear dose response of mCherry to Tat even at early times; (ii) a model where the transactivator is produced as inactive but is then converted to active (i.e., only the transactivator toggles: Tat↔Tat_i_), exhibits a slight threshold in dose response at early times; (iii) a model where both the transactivator and promoter toggle (LTR_OFF_↔LTR_ON_ and Tat↔Tat_i_) extends the transient threshold; (iv) a model where the promoter toggles (LTR_OFF_↔LTR_ON_) and the transactivator toggles between three forms (Tat_ii_↔Tat_i_↔Tat) further extends the transient threshold lifetime. Parameter values used were: {k_α_varied [0.5–10] to generate different Tat-Dendra levels, *k*_*a*_=0.05, *k*_*i*_=0.5, γ=γ2=0.1, *k*_*off*_=0.5, *k*_*on*_=0.1, *k*_*f*_=*k*_*r*_=0.5} and all initial conditions were set to zero (except LTR_OFF_=1).

One prediction of these models (Fig. 5) is that accelerating the promoter toggling transition from *Pr*_*off*_ *to Pr*_*on*_ (increasing *k*_*on*_) should shorten the transient lifetime (for the extreme case: compare *models iii/iv* to *model ii* in Fig. 5 where *k*_*on*_→∞). In support of this, TNF induction increases *k*_*on*_ for the LTR (33, 51), and the data in Fig. 4 show that TNF substantially shortens the lifetime of the transient threshold.

## DISCUSSION

HIV’s ability to establish latency in resting CD4^+^ T lymphocytes remains the chief barrier to curative therapy (22) and an area of active study. Latency establishment does not correlate with viral integration site (23-25) or progressive cellular silencing, and the Tat positive-feedback circuit is necessary and sufficient for latency establishment (26), with epigenetic chromatin silencing possibly maintaining the latent state (31). However, given the non-cooperative nature of Tat feedback (32), the circuit was thought to lack an activation threshold, and so it was unclear how HIV could even temporarily remain in an ‘off’ state to provide an opportunity for the kinetically slower epigenetic silencing mechanisms to act and stabilize latency.

Here, using combination of single-cell analyses (flow cytometry and time-lapse microscopy), we find that the HIV Tat circuit exhibits a *transient* threshold in activation that disappears over time (Fig. 2–3). Promoter activation by TNF shortens the lifetime of this transient threshold (Fig. 4). The transient nature of the threshold accounts for the lack of deterministic bistability and hysteresis in the circuit and previous findings that Tat feedback is non-cooperative (32). We find that a stochastic model, combining two previous models (32, 51), where the transcriptional transactivator and promoter both toggle between ‘active’ and ‘inactive’ states, qualitatively recapitulates the transient-threshold effect (Fig. 5). Other models with additional promoter states (e.g., three-state LTR models) would likely also recapitulate the effect (54).

At its core, the stochastic model generates this threshold—while continuous ODE models do not—because the stochastic model accounts for integer numbers of TF molecules. Thus, the stochastic model intrinsically forms a threshold by requiring a single molecule of acetylated Tat (active TF), rather than a continuous fractional value. Due to rates of conversion, excess molecules are transiently present in the ‘inactive’ state before a single molecule appears in the ‘active’ state, thereby establishing a transient threshold. This effect is interesting to contrast with the other effects of stochasticity in ultrasensitive systems (55).

Physiologically, the transient nature of the threshold may allow the Tat circuit to temporarily remain in an off state and buffer stochastic fluctuations from rapidly triggering positive feedback and active replication, thereby providing a ‘temporal window’ for the kinetically slower epigenetic silencing mechanisms to stabilize the off state. Given the widespread nature of promoter toggling and transcription factor modifications, transient thresholds may be a general feature of inducible promoters.

One caveat to our study is that we only examined a small number of isoclonal integration sites for the LTR promoter. It is possible that these integration sites are somehow unique in their ability to generate a threshold and that higher-throughput analyses of integration sites will produce a different result. It is also important to note that different integration sites yield different effective Hill coefficients (Fig. 2E) and given this range of Hill coefficients, additional integration sites should be analyzed to establish whether the circuit in fact exhibits *H* > 1. If indeed *H > 1,* the model would need to generate a probability of the system being in the active promoter complex [*TF_Pr*_*on*_] that scales with hyperbolic curvature as a function of TF; more formally, there must be some non-zero value of TF where ∂^2^[*P(TF_Pr*_*on*_*)] /* ∂*[TF]*^*2*^ = 0 H.owever, in models *i–iv*, it is straightforward to algebraically show that ∂^2^[*P(TF_Pr*_*on*_*)] /* ∂*[TF]*^*2*^ ≠ 0 for any non-zero value of TF.

To transiently generate *H > 1,* some form of TF cooperativity is required. This cooperativity could in principle be achieved through homo-multimerization (1, 20, 56) of the TF protein, or successive covalent modifications (57) of TF, or successive TF-dependent steps required for promoter activation. However, to recapitulate the data, it is absolutely critical that the mechanism of cooperativity be transient and disappear over time (or disappear as TF levels increase). The homo-multimerization mechanism is the most difficult to reconcile with this. While the active form of Tat might multimerize at early times (low levels of Tat) but then become a monomer at later times (high Tat levels) or under TNF stimulation, this scenario would be an exotic departure from the typical biophysical models of concentration-dependent multimerization of a protein (i.e., monomeric at low concentrations with crowding-induced multimerization). In contrast, it may be more appealing to consider models where at early times (low Tat or LTR-expression levels) two successive Tat dependent steps are required for LTR activation but as the promoter increases in transcriptional activity, one of these Tat-dependent steps becomes a Tat-independent step. For example, active and quiescent promoters differentially localize in the nucleus (58, 59), and if the genomic locus where the LTR integrates repositions as LTR activity increases, the LTR may be subject to different activation signals when it reaches a new nuclear microenvironment (60). In other words, at early times during activation the LTR locus is in a quiescent nuclear micro-environment, whereas at later times after activation, the LTR may reposition to a more “TNF-like” nuclear micro-environment.

There may also exist additional thresholds in LTR activation, such as in response to chromatin remodeling (61). However, as discussed above, the epigenetic chromatin-silencing mechanisms that allow for chromatin-mediated reactivation are dynamically slower effects that cannot explain establishment of latency (26), and thus, this chromatin threshold is likely distinct from the early-time transient thresholding results observed here.

Regarding the potential benefits of such transient thresholding relative to multi-stability, we can only provide speculation. When molecular thresholds are established through self-cooperativity and multistability, it is biochemically difficult to alter the threshold level. In the case of the Tat-LTR circuit, TNF (Fig. 4) and other cellular activators (e.g., trichostatin A, Fig. S6 in Supporting Material) can alter the threshold. Thus, somehow the mechanisms that establish the Tat-LTR threshold are distinct and enable ‘tuning’ of the threshold value. Future work will focus on elucidating the molecular mechanisms that establish the transient threshold and its tunability.

## AUTHOR CONTRIBUTIONS

K.H.A. and L.S.W. designed the research; K.H.A. and E.T. performed the research; K.H.A., E.T., M.T., L.S.W. analyzed data; K.A., M.T., L.S.W. wrote the paper.

## ACKNOWLEDGEMENTS

We thank Brandon Razooky for key initial observations and reagents, and Marielle Cavrois of the Gladstone Institutes Flow Cytometry Core (supported by NIH P30 AI027763, S10 RR028962) and Kurt Thorn of the UCSF Nikon Imaging Center for technical help and advice. K.H.A. was supported by a NSF Graduate Research Fellowship. M.T. acknowledges support from the NIH Office of the Director, the National Cancer Institute, and the National Institute of Dental and Craniofacial Research (NIH DP5 OD012194). L.S.W. acknowledges support from the NIH Director’s New Innovator Award Program (OD006677), and NIH R01 AI109593.

